# Complementary Phenotyping of Maize Root Architecture by Root Pulling Force and X-Ray Computed Tomography

**DOI:** 10.1101/2021.03.03.433776

**Authors:** Mon-Ray Shao, Ni Jiang, Mao Li, Anne Howard, Kevin Lehner, Jack L. Mullen, Shayla L. Gunn, John K. McKay, Christopher N Topp

## Abstract

The root system is critical for the survival of nearly all land plants and a key target for improving abiotic stress tolerance, nutrient accumulation, and yield in crop species. Although many methods of root phenotyping exist, within field studies one of the most popular methods is the extraction and measurement of the upper portion of the root system, known as the root crown, followed by trait quantification based on manual measurements or 2D imaging. However, 2D techniques are inherently limited by the information available from single points of view. Here, we used X-ray computed tomography to generate highly accurate 3D models of maize root crowns and created computational pipelines capable of measuring 71 features from each sample. This approach improves estimates of the genetic contribution to root system architecture, and is refined enough to detect various changes in global root system architecture over developmental time as well as more subtle changes in root distributions as a result of environmental differences. We demonstrate that root pulling force, a high-throughput method of root extraction that provides an estimate of root biomass, is associated with multiple 3D traits from our pipeline. Our combined methodology can therefore be used to calibrate and interpret root pulling force measurements across a range of experimental contexts, or scaled up as a stand-alone approach in large genetic studies of root system architecture.

## INTRODUCTION

In maize, the entirety of primary, seminal, lateral, crown, and brace roots together form a complex architecture which controls the plant’s ability to effectively acquire water, scavenge nutrients, and resist lodging (Hochholdinger, 2009). As a result, root growth and development are fundamental to overall plant development and competitiveness (Lynch, 1995), and several prominent large-effect, loss-of-function mutants in cereal seedling root development have been identified and reviewed previously (Hochholdinger et al., 2017; Bray and Topp, 2018). However, root system architecture of mature, field-grown plants at the quantitative level has been understudied and underutilized due to the relative difficulty in obtaining measurements, with significant tradeoffs intrinsic to any particular phenotyping method (Pauli et al., 2016; Topp et al., 2016). Nevertheless, because root growth is highly plastic and affected by environmental conditions such as substrate moisture and texture (Sharp, 2004; Rich and Watt, 2013), field-based studies are valuable despite their challenges.

In its simplest form, root phenotyping of crop species such as maize or rice can be performed by manual measurement of a limited set of amenable traits, such as root biomass, length, width, or the growing angle, either in soil or soil-free conditions. Currently known genes controlling quantitative root system architecture traits in rice were identified using such measurements, including *PSTOL1* (Gamuyao et al., 2012), *DRO1* (Uga et al., 2013), and a recent *DRO1 homolog* (Kitomi et al., 2020). In field conditions, additional techniques for quantifying roots exist, such as the use of minirhizotrons, soil core sampling, and measuring of root pulling force (Holbert and Koehler, 1924; Bohm and Böhm, 1979; Samson and Sinclair, 1994; Wasson et al., 2014). Historically, root pulling force (RPF) has been useful as a field assay because of its simplicity and potential for scalability, and has been applied to both monocots and dicots (Ortman et al., 1968; O’Toole and Soemartono, 1981; Donovan et al., 1982; Bailey et al., 2002; Landi et al., 2002; Fletcher et al., 2015). RPF is generally correlated with greater root biomass and branching, but more nuanced interpretations and its relationship with recently tractable architectural measurements have yet to be established.

More intricate phenotyping of root system architecture can be performed upon two-dimensional images of either field-excavated root crowns, or young gel-media grown root systems, followed by analysis with specialized software (Le Bot et al., 2010; Lobet et al., 2011; Galkovskyi et al., 2012; Colombi et al., 2015; Das et al., 2015; Symonova et al., 2015; Delory et al., 2018; Seethepalli et al., 2020). Such methods have been used to quantify root system architecture in diverse crops such as maize, wheat, rice, and cowpea (Bucksch et al., 2014; Canè et al., 2014; Burridge et al., 2017; Wedger et al., 2019). However, 2D-based measurements have a limitation in that images are typically taken from only one or two camera perspectives, with information lost from roots occluding each other in the image.

As a result, interest and capacity towards three-dimensional root phenotyping has been increasing, driven in part by technical advances and interdisciplinary approaches (Morris et al., 2017). For example, young cereal plants grown in a gel-based media can be imaged over a 360° rotation, allowing digital reconstruction in 3D and high-throughput feature extraction (Iyer-Pascuzzi et al., 2010; Clark et al., 2011). By scaling this technique to mapping populations, studies have identified new univariate or multivariate root QTLs, demonstrating the value of high-throughput and high-information-content trait capture for dissection of plant architecture (Topp et al., 2013; Zurek et al., 2015). Other 3D-based solutions include the use of X-ray computed tomography (XRT), which is capable of imaging any plant structure, including roots within soil based upon physical density properties (Mairhofer et al., 2012; Mooney et al., 2012; Bao et al., 2014; Rogers et al., 2016; Duncan et al., 2019; Li et al., 2019; Li et al., 2020; Helliwell et al.). While XRT has been applied to plant physiology in some form for nearly two decades, instrument accessibility and technical limitations typically restrict its use to small plant structures, low throughput, and/or limited fields of view.

Here we integrate two protocols, first sampling via RPF and washing mature, lignified, field-grown maize root crowns, followed by imaging via XRT and trait quantification for over 290 roots across multiple field seasons. By imaging the roots absent of soil or other media, scanning and segmentation times were significantly reduced such that replication across two environments and/or two time points was possible. We extracted up to 71 3D features for each root crown sample, including up to 65 traits with significant variation between genotypes, as well as root shape or distributional traits, which showed differences between experimental contexts. The median broad-sense heritability across all traits ranged from 0.23 to 0.56, depending on the germplasm and conditions. Finally, we examined covariance between 3D traits and RPF values to identify correlations between high-resolution phenomics and high-throughput field data. This study illustrates the contributions of both phenotyping approaches, and provides insights into the root architectural attributes that influence RPF, which can be used for the mapping of root traits, multi-environment studies, and crop breeding.

## MATERIALS AND METHODS

### Plant Germplasm and Experimental Design

All plants were grown at the Colorado State University Agricultural Research Development & Education Center in Fort Collins, CO, USA (40.649 N, −105.000 W) in 2017 and 2018. The *Genomes 2 Fields* (G2F) germplasm (https://www.genomes2fields.org/) was planted in May 2017 in a split-plot design with full irrigation or limited irrigation (drought) treatments, with two field replicates per treatment for a total of 1060 plots total. Prior to planting, the field was fertilized with nitrogen at 65 lbs per acre. From 260 genotypes planted as part of a large field trial, 30 genotypes were selected for root imaging in this study across both irrigation treatments (Supplemental Table 1), for a total sample size of *N* = 107 roots.

The Shoot Apical Meristem (SAM) diversity panel (Leiboff et al., 2015) along with 11 hybrid and 4 inbred check lines were planted in May 2018 using a split-plot design with full irrigation or limited irrigation (drought) treatments, with three field replicates per treatment. Prior to planting, the field was fertilized with nitrogen at 190 lb per acre. From 390 genotypes planted as part of a large field trial, 20 genotypes were subsampled for root imaging in this study across both irrigation treatments (Supplemental Table 1). Root systems were harvested at 9 weeks after planting (time point 1) and again at 16 weeks after planting (time point 2), for a total sample size of *N* = 187 roots.

In both the G2F and SAM experiments, each plot consisted of two 12-foot rows with 30-inch spacing between rows and 9-inch spacing between plants within rows. The irrigated treatments received approximately 1 inch of water per week, while the drought treatments were irrigated until well established (approx. 5 weeks after planting) and then received only natural precipitation (103.8 mm and 69.9 mm in the 2017 and 2018 growing seasons, respectively), except at the root harvesting when it also received irrigation to homogenize the root harvesting process.

### Field Phenotyping and 2D Root Imaging

The protocol used for root pulling and harvesting was similar to that in (Fletcher et al., 2015). Briefly, all plants were irrigated 24 hours prior to sampling to homogenize soil conditions at root harvest. Maize plants were tied at the base of the stem, just above the root crown, with a rope attached to a dynamometer. The root system was extracted from the soil by vertical manual pulling, with the required force (Kg) needed measured using a hand-held Imada DS2 digital force gauge (Imada Inc., Northbrook, IL, USA). Within each field treatment (full vs limited irrigation), two roots per genotype were harvested from the G2F population and an average of 4 roots per genotype (across two time points) were harvested for the SAM population. After pulling, root samples were washed to remove all remaining soil and allowed to dry before imaging.

Roots from the G2F 2017 experiment were also imaged in 2D using a photography station equipped with a Sony a7 II mirrorless camera. Roots were placed horizontally on a flat surface with a black background, and the resulting images were then cropped and analyzed using DIRT (Das et al., 2015).

### 3D Root Imaging and Feature Extraction

For 3D phenotyping in the G2F and SAM populations, root samples were clamped at the stem with a small vise and imaged using a North Star X5000 X-ray system (North Star Imaging, MN, USA). The sample was continuously imaged while rotated using the North Star efX-CT software system, generating 1800 radiographs per sample (approx. 3 minutes). To provide an internal calibration of the image geometry, a fixed standard (15 mm large tool, North Star Imaging) was imaged with each sample batch. The radiographs were then reconstructed using efX-CT and exported as an unadjusted RAW volume, resulting in a voxel size of 109-113 μm depending on the sample batch. For each sample, the RAW volume was converted to 2D slices using the custom Python script *raw2img*. The slices were then thresholded and binarized using the custom script *batch-segmentation*, which subsequently converted each sample to a 3D point cloud and quantified 19 root traits adapted from (Galkovskyi et al., 2012). Finally, each sample was converted to a 3D skeleton using the custom script *batch-skeleton*, which then quantified an additional root 52 traits. The raw phenotype data is available in Supplemental File 1.

A list and description of root features measured using *batch-segmentation* and *batch-skeleton* are available in Supplemental Table 2. Mean, standard deviation, skewness, kurtosis, energy, entropy, and smoothness from the distributions of root biomass (volume), convex hull, and solidity were calculated using the method described in Malik and Baharudin, 2012. Fractal dimension, which measures the degree to which root subsections approximate a smaller copy of the whole root crown, was estimated by taking the 2D projection of the 3D volume, then calculated using a similar approach to that described in Grift et al., 2011. DensityS features are computationally similar to plant compactness traits described in Yang et al., 2014. Scripts used for image processing and feature extraction are available at https://github.com/Topp-Roots-Lab/

### Statistical Analysis

All downstream (i.e. post feature extraction) analysis was performed in the R statistical computing environment. Initially, principal component analysis using all 71 3D roots traits was used to identify large outliers, leading to the removal of 2 samples in the G2F 2017 data and 3 samples in the SAM 2018 data. Additionally, for all univariate analysis, outliers within each trait were identified and omitted if they were beyond the 1st quartile minus 1.5 * interquartile-range or the 3rd quartile plus 1.5 * interquartile-range.

After univariate outlier removal, analysis of variance (ANOVA) was performed for each trait using the *car* package (Fox and Weisberg, 2018). Subsequently, the ANOVA p-values were adjusted using the Benjamini-Hochberg method. Individual two-sample comparisons as seen in boxplots were performed using Mann-Whitney U tests. Correlations between root traits were calculated using Spearman’s correlation coefficient. Linear regressions were performed using the *lm* function in R.

Variance components were estimated by using the *lme4* package (Bates et al., 2015) to fit the linear model *Y_ijk_ ~ G_i_ + E_j_* + (*G*E*)_*ij*_ + *e_ijk_*, where *Y* is the phenotypic value, *G_i_* is the *i*^th^ genotype, *E_j_* is the *j*^th^ environment, (*G*E*)_*ij*_ is the interaction between the *i*^th^ genotype and the *j*^th^ environment, and *e_ijk_* is the residual error of the *k*^th^ sample from the *i*^th^ genotype and *j*^th^ environment. Broad-sense heritability was calculated using the equation 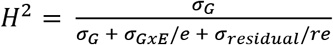 Where σ_G_ is the estimated phenotypic variance due to genotype, σ_GxE_ is the estimated phenotypic variance due to genotype x environment, *σ_residual_* is the residual variance, *e* is the number of environments, and *re* is the average number of biological replicates per genotype across both environments (Nyquist and Baker, 1991). This heritability estimator is optimized as a predictor of the response to selection. For the SAM 2018 data, variance components and broad-sense heritability were calculated separately for the two time points.

Principal component analysis (PCA) of the 3D root data within the G2M and SAM experiments was performed using the base R *prcomp* function. For PCA-LDA, PCA was performed upon each genotype subset, and the number of principal components required to explain 90% of the trait variance was used as inputs into the LDA function from the *MASS* package (Venables and Ripley, 2002).

The *randomForest* (Liaw et al., 2002) and *caret* (Kuhn and Others, 2008) R packages were used for random forest classification, with *mtry* and *ntree* parameters found using a grid search approach between every combination of *mtry* between 1 to 20 and *ntree* values of 500, 1000, 2500, 5000. Parameters giving the best accuracy were kept, as calculated by 10-fold cross validation repeated 3 times. From the final random forest models, the proximity matrix was calculated and non-metric multidimensional scaling was used to visualize the distances between samples.

## RESULTS

### Field and 3D Phenotyping Capture Variation in Maize Root System Architecture

In each of two field seasons, we sampled 50 maize genotypes (30 from the G2F panel and 20 from the SAM panel) that were grown under two different irrigation treatments, providing two environments in terms of soil moisture. At the designated time point(s) for sampling (see Methods), root crowns were excavated by tying the base of the stem with rope to a digital force gauge, and manually placing a vertical force on the plant until the root crown was ruptured and lifted free from the soil. The force gauge attached to the root system measured the kilogram of force required for this process, also known as the root pulling force (RPF).

Field-pulled root samples, which maintain their 3D structure due to lignification, were washed clean and subsequently imaged using a Northstar X5000 X-Ray computed tomography system (Figure 1, Supplemental Figure 1A). Scans were then exported as vertical (y-axis) image slices, thresholded using an automated algorithm, and converted to a skeletonized image and a point cloud image for trait analysis (Supplemental Figure 1B-D). These were analyzed for 19 3D traits using an established 3D-analysis pipeline (Bray and Topp, 2018) as well as 52 additional traits predominantly focused on 3D root distribution traits using a newly developed series of feature extraction tools. In total, 71 traits from the 3D volumes were extracted and used for analysis (Supplemental Table 2).

**Figure 1:**
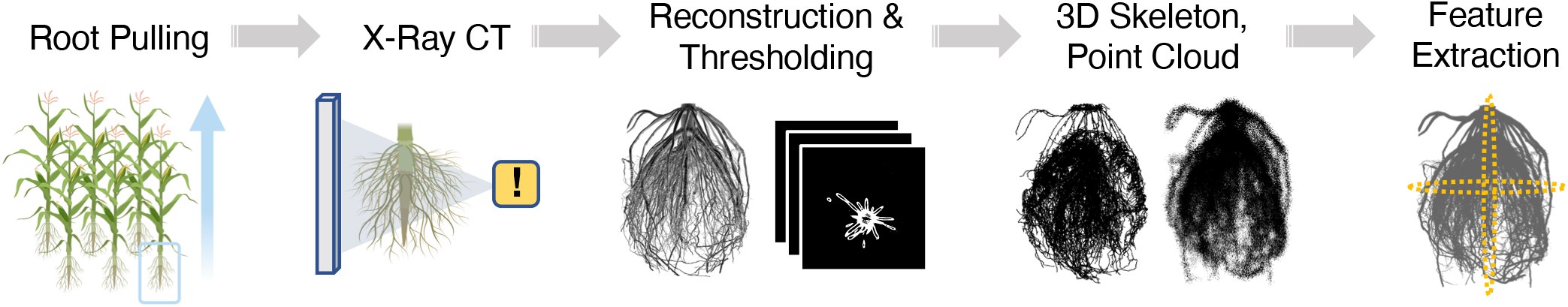
Pipeline for 3D root imaging using X-ray computed tomography. Samples are excised from the soil in the field using the root pulling force method, and the measurement recorded. The root crown is washed, dried, and then imaged using the NorthStar Imaging X5000 (see Methods) to generate radiographs as the sample is rotated 360° across the vertical axis. From the radiographs, a 3D reconstruction is generated using the FDK algorithm. Slices along the vertical axis are exported for automated thresholding, from which a skeleton and point cloud model of the root crown are generated. 3D root traits are then measured from the skeleton and point cloud, and analyzed. See Supplementary Figure 1 for large example images of 3D models.

In the G2F data set, root samples were also photographed for 2D image analysis via DIRT using recommended protocols (Supplemental Figure 1E) (Das et al., 2015). Correlations between the most directly comparable 2D and 3D traits were as we expected - for example, 2D area and 3D surface area had a Pearson correlation coefficient of 0.754.

To assess the degree to which traits derived from field-pulled root crown samples would respond to selection, we estimated broad-sense heritability (*H^2^*) in the G2F experiment for each 3D and 2D trait, as well as for RPF (Supplemental Figure 2). Traits related to overall root crown size showed similar *H^2^* values between 3D measurements (e.g. 3D surface area *H^2^* = 0.47) and 2D measurements (e.g. 2D area *H^2^* = 0.44). The traits with highest heritability, however, were 3D-derived maximum root count (*H^2^* = 0.76) and average root radius (*H^2^* = 0.74), illustrating where 3D root phenotyping is particularly adept. Among 2D traits, high heritability did not necessarily result in high association with RPF, although some traits such as 2D area had a strong positive association (Supplemental Figure 2E-F). Nevertheless, depending on the experimental conditions and amount of replication, it is probable that many root traits - though computationally extractable - have questionable value due to high background noise and sensitivity to sampling variation. In total, for example, 19 3D traits and 33 2D traits had calculated *H^2^* values of less than 0.05; therefore, both 3D and 2D root traits must be screened and evaluated for a given data set before drawing conclusions. Reassuringly, however, RPF itself had a *H^2^* value of 0.67 in the G2F experiment, which is remarkably high for a physical field-based root measurement, even when compared to the architectural traits measured from 3D images.

In the SAM experiment, broad-sense heritability for 3D traits were calculated separately within each time point (Supplemental Figure 3A-C). In general, *H^2^* values here were higher than in the G2F experiment, which reflects a combination of the field conditions, genetic variation, and sample sizes. As an average across both time points, root crown width (“HorEqDiameter”, mean *H^2^* = 0.81), fractal dimension top view (mean *H^2^* = 0.81), maximum root count (mean *H^2^* = 0.80), convex hull volume (mean *H^2^* = 0.79), and surface area (mean *H^2^* = 0.79) were among the most heritable traits, although the individual performance of these traits fluctuated depending on the time point. However, the average heritability across all 3D traits was only slightly higher at first time point (0.510) than at second time point (0.505), suggesting that heritability of most roots traits is relatively static over this time span. One interesting exception to this is RPF itself, which had a *H^2^* value of only 0.59 at the first time point, but significantly increased to a *H^2^* value of 0.85 by the second time point. This indicates that RPF measurements taken later in the plant life cycle may be more informative and reliable for the purposes of distinguishing genotypic differences in maize root system architecture, as well as for breeding. Overall, however, traits with higher average heritability across time points tended to also have a greater correlation in measurements between time points (Supplemental Figure 3D).

Focusing on 3D root traits and RPF, we subsequently wanted to examine whether genotype and environment effects were significant factors on a trait-by-trait basis (Figure 2). Using analysis of variance (ANOVA), in the G2F experiment we detected 21 root traits where genotype had a significant effect, and 35 traits where the environment (irrigation regime) had a significant effect (Figure 2B, Supplemental Figure 4A, Supplemental Figure 5). Root traits affected by both genotype and environment include RPF, average root radius, median/maximum number of roots, convex hull skewness, and solidity in several regions along the middle of the root crown. Non-parametric tests for differences between environments confirmed that RPF, average root radius, and convex hull volume, for example, were higher in the high irrigation environment, whereas solidity was higher in the low irrigation environment (Figure 2C). These meet expectations of soil moisture effects on root system architecture (e.g. more expansive growth under higher moisture availability), providing confidence to our 3D phenotyping.

**Figure 2:**
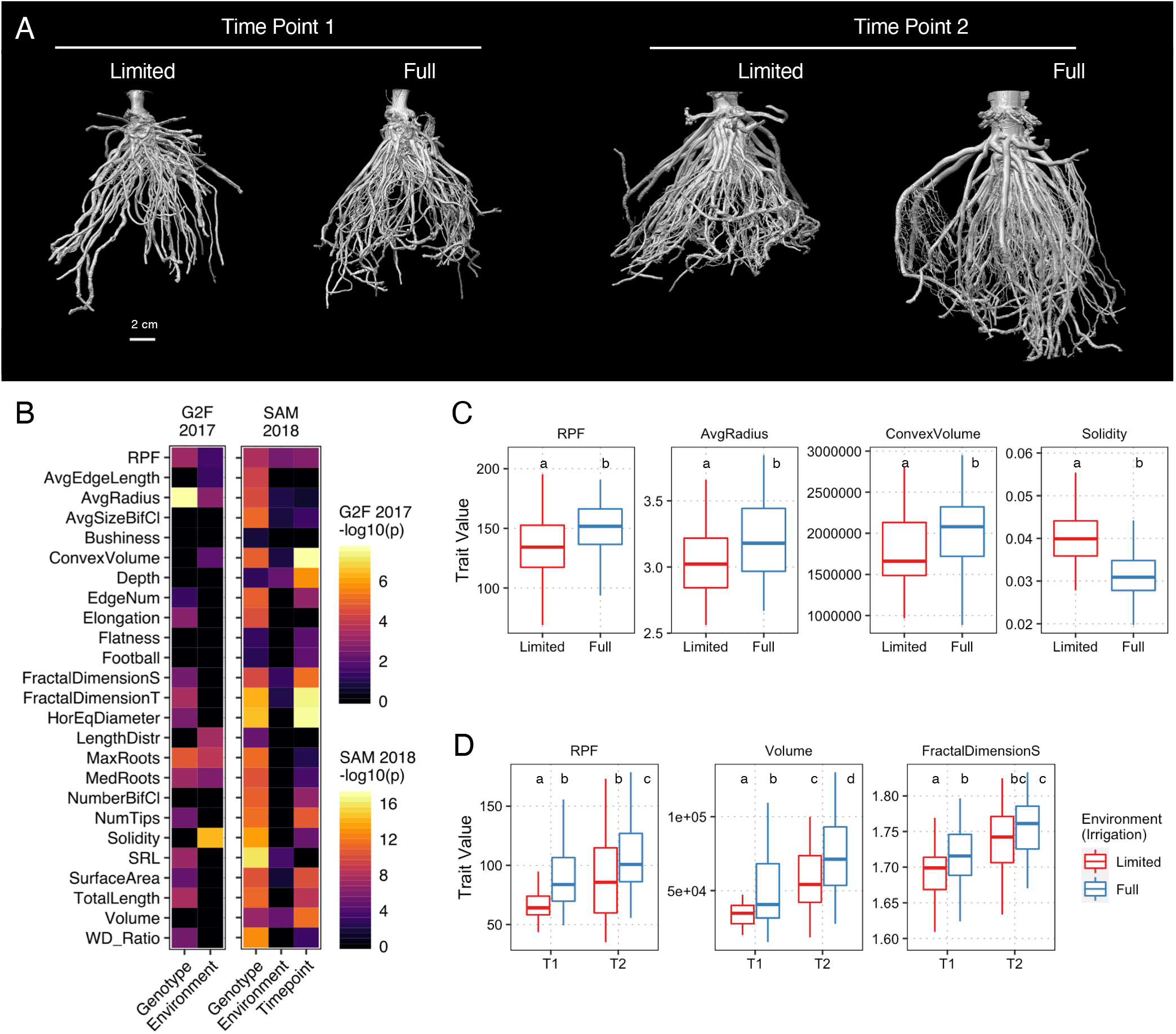
RPF and 3D global root system architecture traits are affected by genotype, environment, and developmental time point. (A) 3D reconstructions from X-ray imaging of genotype Tx601 root crowns in the SAM 2018 experiment, at the two time points (9 vs 16 weeks) and from each environment (limited vs full irrigation). (B) ANOVA for genotype, environment, and time point (in SAM 2018) effects upon RPF and 3D root traits (adjusted p < 0.05); for legibility, G2F 2017 and SAM 2018 experiments were separately scaled, and non-significant features were set at a -log10(p) value of 0. (C) Boxplot of selected traits significantly different between the two environmental conditions in the G2F 2017 experiment (Mann-Whitney U Test p < 0.05). (D) Boxplot of selected traits significantly different between the two development time points and/or the two environmental conditions in the SAM 2018 experiment (Mann-Whitney U Test p < 0.05).

In the SAM experiment, the situation was somewhat reversed: in support of the overall higher trait heritability, a remarkable 65 root traits had a significant effect from genotype, but only 17 traits had a significant effect from environment, while 37 traits had a significant effect from time point (Figure 2B, Supplemental Figure 4A, Supplemental Figure 6). Traits such as RPF, surface area, volume, root crown depth, fractal dimension side view, and biomass distribution skewness were affected by all three variables. Again, non-parametric tests for differences in RPF, volume, and fractal dimension side, for example, confirmed the impacts of environment and time point as detected by ANOVA (Figure 2D). The somewhat divergent trends in 3D root phenotypes between the G2F and SAM experiments, however, indicate that additional generalizations about root system architecture and how growth plasticity relates to it may be difficult to come by, as root variation is highly dependent on the experimental conditions and/or genotypes. Indeed, although the sample sizes here precluded strong statistical power to test genotype-environment interactions using ANOVA, variance component analysis suggests that such interactions may have a significant influence on a number of root architecture traits (Supplemental Figure 2B; Supplemental Figure 3B-C).

Next, we applied supervised multivariate classification methods to determine which traits were most closely associated with differences in genotype, environment, or time (Supplemental Table 3). Because of the high number of genotypes (18 in the G2F set and 16 in the SAM set, after filtering for genotypes with the least missing data), in both experiments the data was split into every possible combination of three genotypes, generating 816 different genotype combinations in the G2F set and 560 different genotype combinations in the SAM set. We performed PCA-LDA for genotype classification upon each three-genotype data subset, in each case using the minimum number of principal components to explain 90% of the variance (5-7 PC’s with a median of 6 in G2F data; 9-13 PC’s with a median of 11 in SAM data) as the inputs for LDA (Figure 3A-B). Across all genotype combination subsets, the average classification accuracy using leave-one-out cross-validation was 54.6% in the G2F and 67.2% in the SAM, both significantly higher than the 1/3 expected by random chance, particularly considering the realistic possibility that numerous genotypes may in actuality be phenotypically similar.

**Figure 3:**
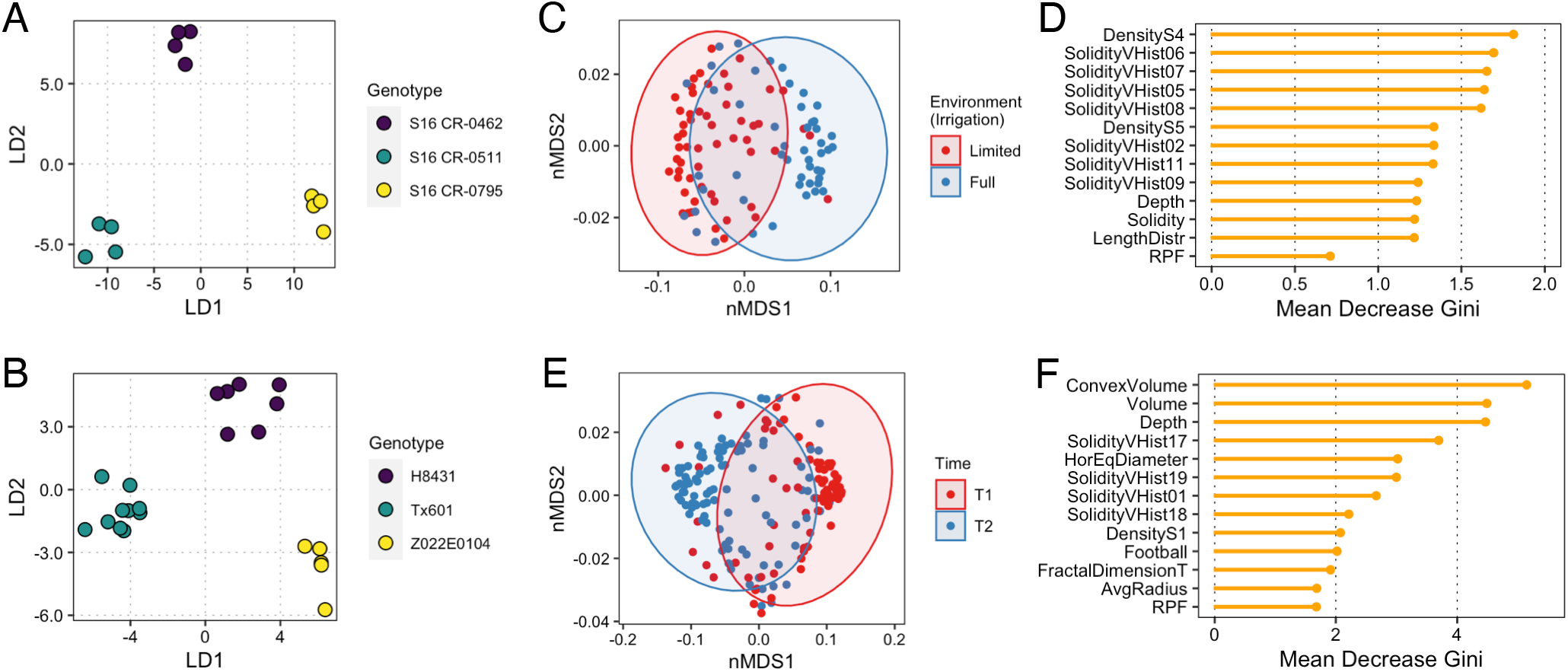
Classification based on genotype, environment, and time point using 3D root system architecture traits and RPF. Examples of highly distinguishable genotypes by PCA-LDA in the G2F 2017 data (A) and SAM 2018 data (B). Random forest classification of all samples based on environment in the G2F 2017 data (C) and importance of the 12 most influential traits plus RPF (D). Random forest classification of all samples based on time point in the SAM 2018 data (E) and importance of the 12 most influential traits plus RPF (F).

We examined what traits were most important towards PCA-LDA classification across all genotype combinations (Supplemental Figure 7-8). In both the G2F and SAM populations, maximum root count, average root radius, and specific root length tended to be very important for genotype discrimination. Additionally, the median root count, number of root tips, elongation, and average edge length were important in genotypic classification among the G2F population, while several root solidity and density traits were important in genotypic classification among the SAM population. Although RPF was not among the top traits for genotypic classification, it was still well above average, ranking 12th overall in the G2F set and 19th overall in the SAM set. Interestingly, traits related to overall root size such as volume or surface area did not seem to be important factors overall in discriminating between genotypes, suggesting that these metrics, although intuitive and undoubtedly important in other contexts, are by themselves insufficient to distinguish between multiple and often subtly distinct genotypes, highlighting the need for the more comprehensive phenotyping described here.

To evaluate the overall effect of the environment (influential in the G2F experiment) and time (highly influential in the SAM experiment), we performed random forest classification to distinguish between the two possible levels of each variable upon root system architecture. For these classifications we included all genotypes, which increased the sample size for each model. Using 10-fold cross validation, the best model parameters resulted in a classification accuracy of 81.0%, indicating that while the environment had an effect which was detectable using classification techniques, the contrasting irrigation regimes were not so dramatic as to result in a shift in root system architecture obvious across every sample (Figure 3C). Nevertheless, changes in density and solidity distributions, as well as root crown depth, were the most distinguishing features, with RPF being less important (Figure 3D). Here, the importance of solidity distributions in the upper half of the root crown (including SolidityVHist 02-11) is consistent with ANOVA analysis (Supplemental Figure 4A); in particular, the low broad-sense heritability of SolidityVHist 05-10, coupled with disproportionately high variance from environment and genotype-environment effects, indicates that these are more determined by environmental factors than by genetics in this experiment (Supplemental Figure 2A-B). On the other hand, DensityS5 (a measure of relative compactness), which had a moderately high heritability in this experiment, is still important for distinguishing the effect of environment, suggesting that this trait is strongly affected by both genotype and environment.

For classifying roots based on time point in the SAM data, using 10-fold cross validation the best model parameters resulted in a random forest classification accuracy of 78.6% (Figure 3E), which was reasonable when considering that samples across both environmental conditions were included. Here, differences in convex hull volume, volume, depth, root crown width, and solidity distribution were the most distinguishing features, with RPF closely behind these and other important traits (Figure 3F). Furthermore, solidity distribution features at the very top and bottom of the root crown (SolidityVHist 01 and 17-19) appear to be relevant. Again, these results are unsurprising given the expectation of increasing root crown size over time, and are largely consistent with ANOVA results on time point effects upon these traits (Figure 2B).

### Root Architecture Relationships and Correspondence to Root Pulling Force

Root pulling force has been used historically and recently as a proxy for root biomass and root volume. Nevertheless, to have additional and more detailed information on the architectural changes that RPF measures would increase its utility as a field assay. We first calculated correlations between RPF and 3D root phenotype across all measured samples, irrespective of genotype and environment, or time point in the case of the SAM data. In both experiments, RPF was most correlated with root volume, fractal dimension, surface area, total root length, root crown width, number of bifurcating clusters, and number of root tips. (Figure 4A-B, Supplemental Table 4). Traits negatively correlated with RPF were generally weaker and less consistent between the two experiments, but did include convex hull energy (a measure of root system uniformity) and Density S5 in both cases.

**Figure 4:**
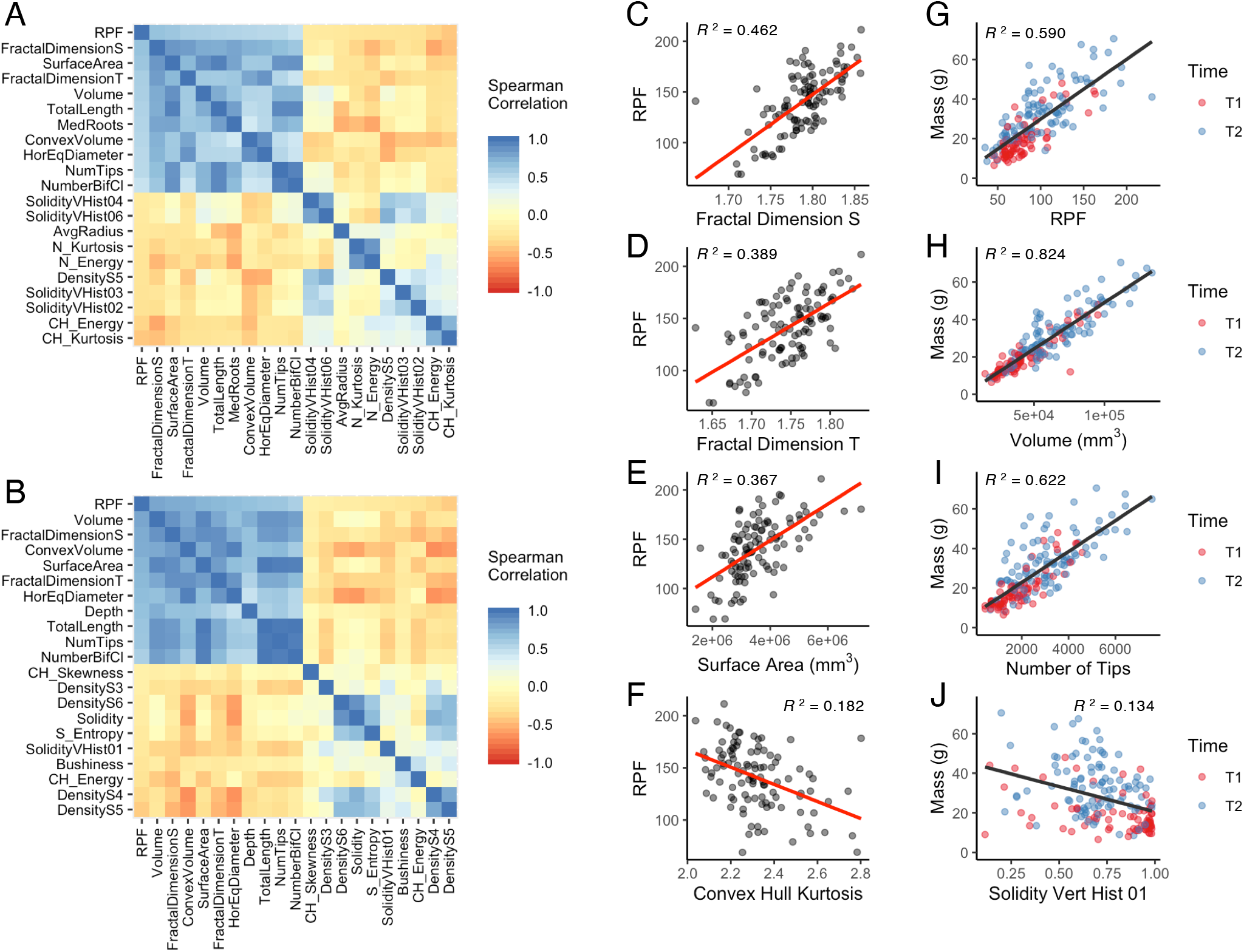
RPF and 3D root system architecture traits are strongly associated with each other and root biomass. Heatmap of RPF and its 20 most correlated traits (10 most positive and 10 most negative) in the G2F 2017 (A) and SAM 2018 (B) datasets. Regression of example traits positively correlated to RPF such as fractal dimension side/top (C, D) and surface area (E), and example traits negatively correlated to RPF such as convex hull volume (F), within the G2F 2017 dataset. Regression of example traits positively correlated to root biomass such as RPF (G), volume (H), and number of tips (I), and example traits negatively correlated to root biomass such as solidity vertical histogram-01 (J), within the SAM 2018 dataset. Adjusted r-squared values for C-J shown in respective plot insets.

Associations between root pulling force and root system architecture traits are most useful if they are not only significantly correlated, but also exhibit a close linear relationship. Regression analysis between RPF and positively correlated 3D architecture traits (as observed in the G2F experiment), such as fractal dimension and surface area, showed a reasonably good fit (Figure 4C-E). In contrast, there was a relatively poor fit with convex hull kurtosis, the most negatively correlated trait (Figure 4F). These may in part be due to sampling error or noise, but also because multiple root characteristics that may not be strongly correlated to each other nevertheless each contribute to RPF in various ways. Nevertheless, the regression fit between physical root biomass (i.e. root crown weight) in the SAM experiment and RPF or other positively correlated 3D architecture traits was extremely high, while again relatively poor with negatively correlated traits such as solidity in the upper root crown (Figure 4G-J). This high goodness-of-fit was not a by-product of regression between two time points; rather, regression between physical root biomass and these 3D traits remained high even when observing trends and regressions within each time point (Supplemental Figure 4B-J). Furthermore, regression fit was typically higher in time point 1 than in time point 2, which might be due to root crown traits beginning to diverge in ways more independent of biomass, such as in architectural and spatial orientation, which could nonetheless contribute to RPF.

To explore the degree to which trends across multiple traits may be associated with RPF, we performed principal component analysis (PCA) from the G2F and SAM data using the 3D-based root phenotypes alone (Supplemental Figure 9A, E). In the G2F data, RPF was more tightly associated with principal component 2 (Supplemental Figure 9B), which was primarily composed of traits related to overall size, i.e. surface area, volume, total root length, and number of root tips, but also significantly composed of 3D biomass distribution traits (Supplemental Figure 9I). On the other hand, in the SAM data, RPF was more tightly associated with principal component 1 (Supplemental Figure 9F), which as with the G2F data was primarily composed of traits related to overall size, including surface area, volume, and total root length, and additionally fractional dimension side/top, but notably not of 3D biomass distribution traits (Supplemental Figure 9J). Both PC1 and PC2 were statistically different between the two environmental conditions in the G2F data and between the two time points in the SAM data (Supplemental Figure 9C-H), but the differences between the G2F and SAM results here likely derive from the fact that much of the phenotypic variation in the SAM data is greatly affected by sampling time point, which an unsupervised method such as PCA does not distinguish.

As a whole, multiple analytical approaches corroborate the conclusion that distinct sets of root traits are relevant depending on the germplasm, environment, and developmental time stage, reinforcing the relevance of high-dimensional root phenotyping. This also demonstrates not only the complexity of root architectures, but its propensity to change under different contexts and genetic influences, and its ability to adapt to various conditions.

## DISCUSSION

We have shown that using X-ray computed tomography, changes in 3D root architecture between different treatments and conditions in the field can be measured in a biologically interpretable way, while also with high precision and detail. For example, soil moisture conditions affect the solidity of the root system, particularly in the mid-portion of the crown. In contrast, changes over time influence not only the overall size of the root crown, but also solidity in the upper and lower portions of the crown. Interesting, in both contexts, the depth (i.e., the length of the vertical axis) of the root crown also was a distinguishing feature. This is likely specific to root crowns excavated using the root pulling method, as under standard shovel excavation, the root depth would be arbitrary. By using root pulling, however, the depth is a function of the root system and the soil conditions, as these determine where the root crown breaks and therefore encapsulates some information. Finally, in many instances the 3D architectural measurements were easily sufficient to distinguish different maize genotypes, primarily using an entirely different set of traits including specific root length, median/maximum root count, average root radius, and so on.

To date, 2D imaging has been by far the most popular form of quantifying root system architecture, whether in *Arabidopsis* grown on media plates, or root crowns excavated from the field. While relatively straightforward and convenient, 2D imaging does not represent true root system architecture in its natural form, and therefore may omit important information. More recently, optical imaging platforms have been developed to perform 3D imaging of plants growing in gel media (Clark et al., 2011; Topp et al., 2013; Jiang et al., 2019). Here, we present a new approach to quantifying hundreds of field-excavated root crowns using X-ray CT, which is typically restricted to very small sample sizes or reconstituted soils from pot experiments (Bao et al., 2014). Our results suggest that 3D imaging and the root architectural traits derived from it have higher heritability, and therefore may be more informative, than methods using 2D imaging. Therefore, we anticipate that future studies and breeding efforts in quantitative root system architecture will increasingly utilize 3D phenotyping.

Nonetheless, the significant overhead associated with 3D imaging and analysis of roots will be a limitation to many researchers for the foreseeable future. We addressed this by making explicit comparisons between high information content 3D phenotypes and root pulling force, which is accessible and can be scaled to high throughput levels. RPF measurements had several significant positive correlations with 3D architecture traits including volume, surface area, total root length, number of roots, and fraction dimension. Indeed, fractal dimension was a surprisingly powerful trait, not only being highly correlated with RPF, but also having high heritability and contributing significantly to differentiating root system architecture over time. This is additional evidence that fractal dimension of root systems (Tatsumi et al., 1989; Eghball et al., 1993; Nielsen et al., 1997; Eshel, 1998; Grift et al., 2011), is indeed a useful feature for quantifying maize root crowns under a variety of scenarios.

However, there remains sufficient sources of variance among features such that future improvements could strengthen the relationship between the above traits and RPF values. For example, we performed root pulling manually by hand, but a field robot or other form of mechanical assistance may result in more consistent RPF measurements (Mayer, 2019). Furthermore, while the fields were flooded just prior to root pulling to standardize the soil moisture conditions at the time of RPF sampling, local heterogeneity in soil texture or compactness may influence the measurements. This can be addressed in part by integrating larger studies whereby spatial effects can be modeled, which likewise would be facilitated with mechanization of the root pulling process. Finally, it should be noted that several traits not measurable by our XRT system could contribute to RPF, including the abundance of root hairs and variation in rhizosheath formation. Indeed, the fact that the heritability of RPF here increased later in development suggests that it may be controlled by different genetic factors over time, and therefore it may also be possible to select and breed for early and later root phenotypes at least partially independently.

A better understanding of the exact nature and the potential interactions between the many root traits investigated here, such as exactly how fractal dimension and root volume together affect RPF, as well as the addition of more traits that could theoretically be calculated from 3D imaging (such as those relating to topology), may lead to additional insight into the relationship between root system architecture and RPF. Fully resolving this relationship would be particularly beneficial for multi-environment phenotyping, which requires high sample sizes to which the RPF method is well-suited for. Indeed, our study suggests that many more field-scale studies, utilizing wide-ranging conditions and germplasms, will be needed to fully characterize and understand quantitative root system architecture and genotype-environment interactions in diverse plant species such as maize.

