## Supplemental Figures for "Complementary Phenotyping of Maize Root Architecture by Root Pulling Force and X-Ray Computed Tomography"

**Supplemental Figure 1:** Example root crown from genotype S16 CR-0511 in the G2F 2017 experiment showing the X-ray CT reconstructed 3D volume (A), point cloud (B), close-up of a region in the point cloud (C), skeleton (D), and an independent 2D image (E).

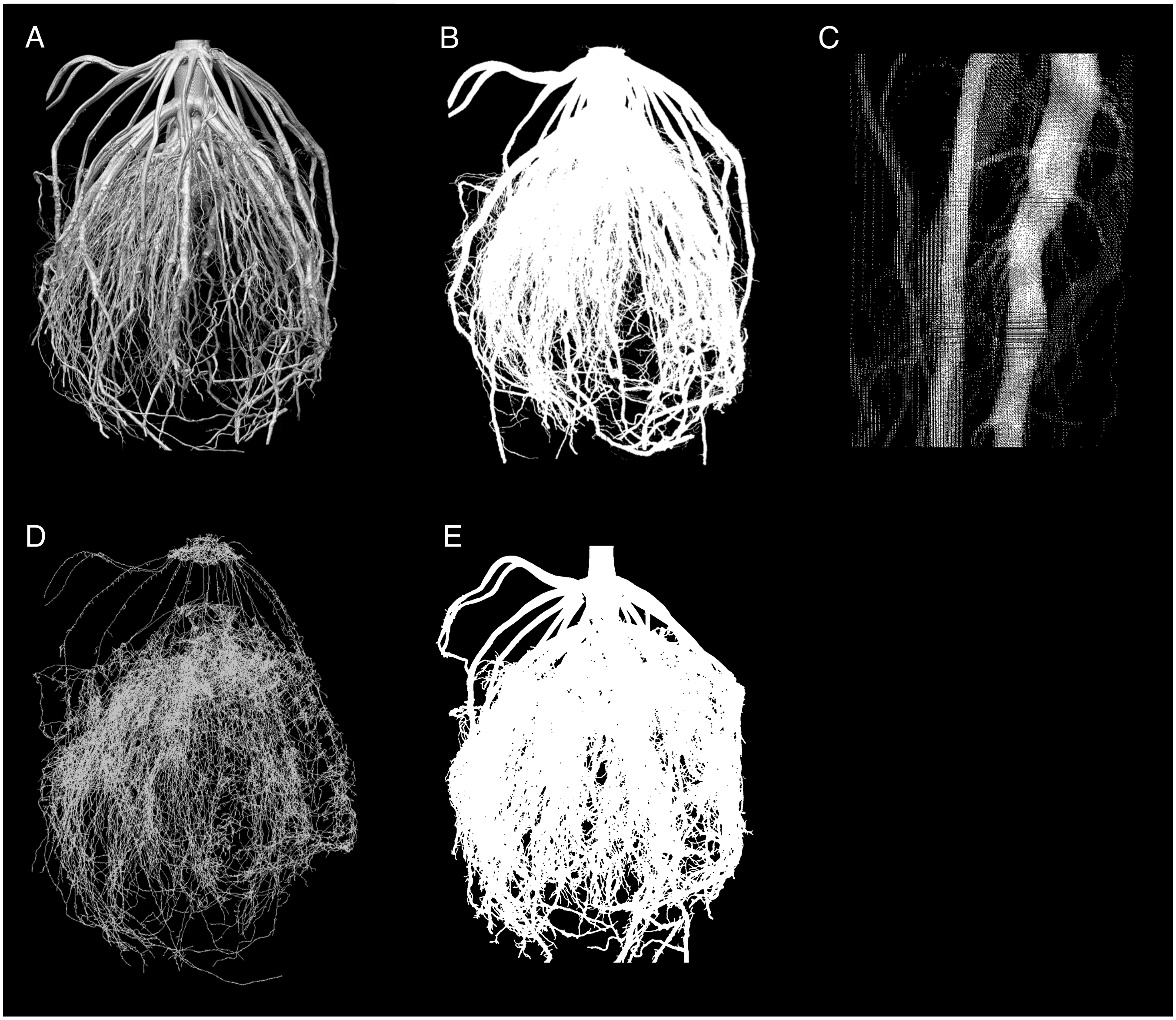

**Supplemental Figure 2:** Broad-sense heritability (A) and variance component analysis (B) of RPF plus all 3D root traits from the G2M 2017 experiment. Broad-sense heritability (C) and variance component analysis (D) of 2D roots traits with heritability > 0 from the G2M 2017 experiment. Linear regression between RPF and 2D root area (E) or NR_RTP_SEG_II (F).

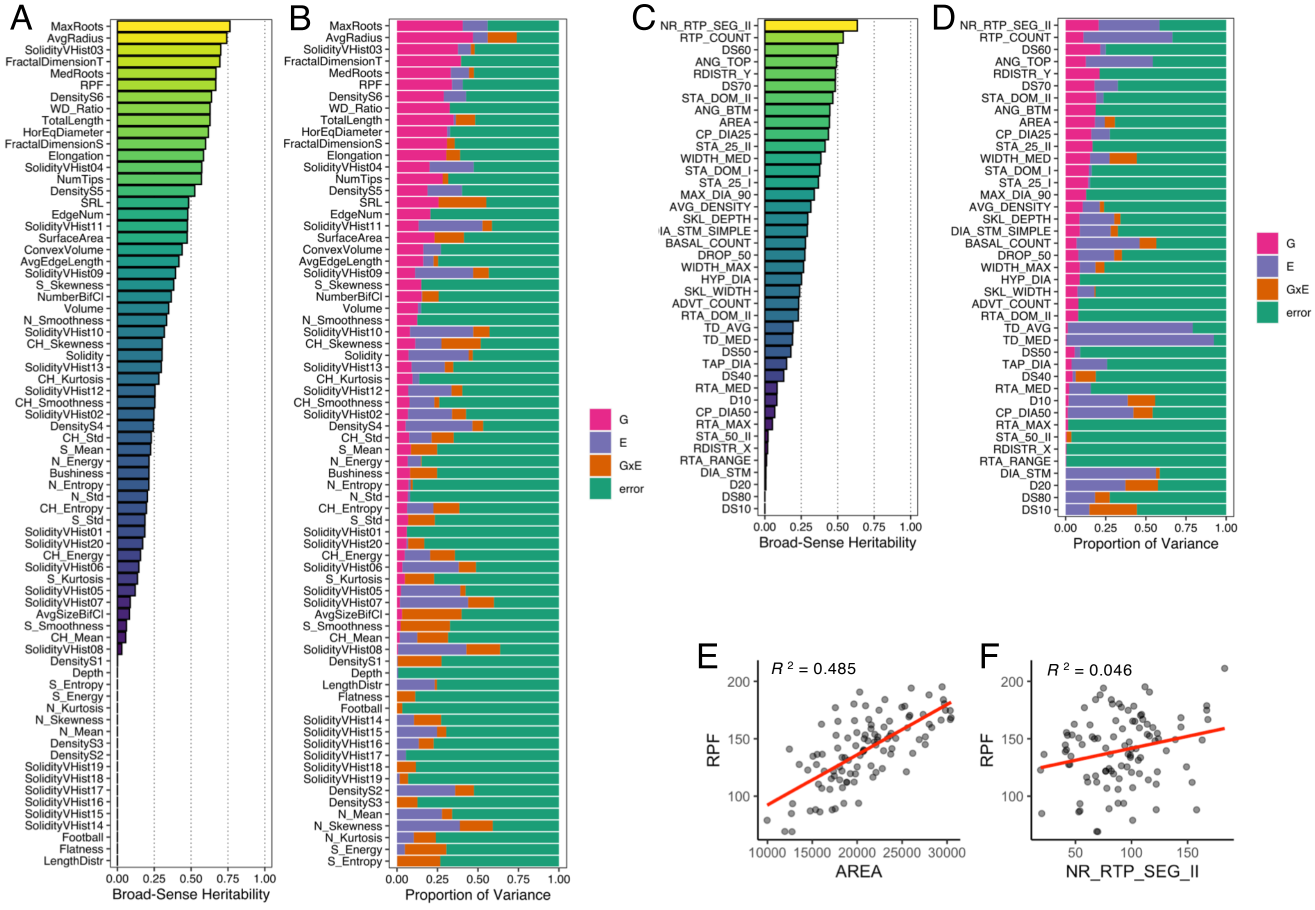

**Supplemental Figure 3:** Broad-sense heritability of RPF and all 3D root traits from the SAM 2018 experiment at both time points (A). Variance components for each trait in time point 1 (B) and time point 2 (C). Traits in A-C are ordered by mean heritability across both time points. Scatterplot of mean heritability across both time points versus Spearman correlation between both time points, with each trait as a single point (D); correlation values for all traits are contained in Supplemental Table 5.

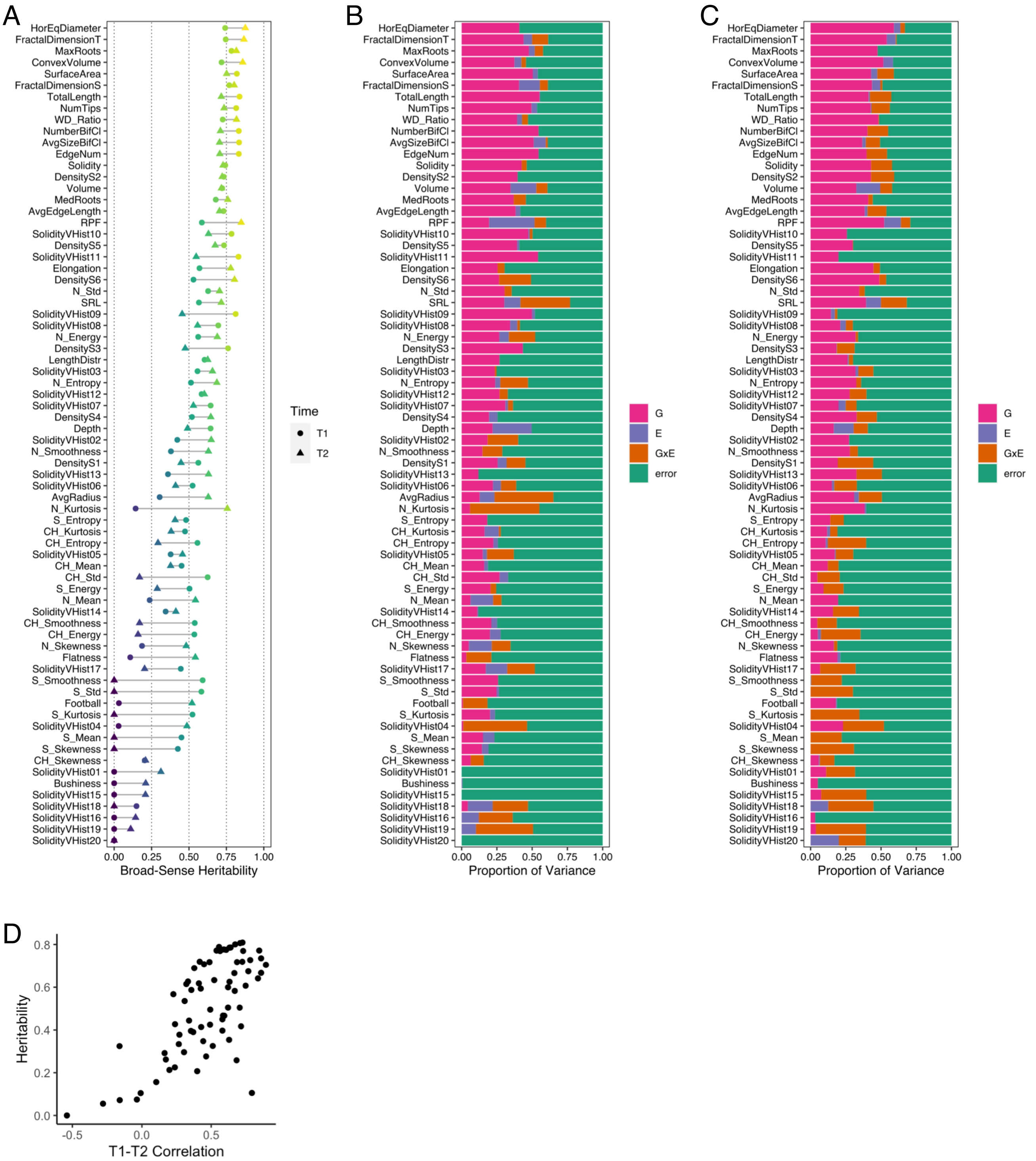

**Supplemental Figure 4:** RPF and 3D distributional root system architecture traits are affected by genotype, environment, and developmental time point. There are consistent relationships between root biomass and RPF or 3D root traits across both time points (B-H), and a non-linear relationship between fractal dimension side/top and root biomass (I-J).

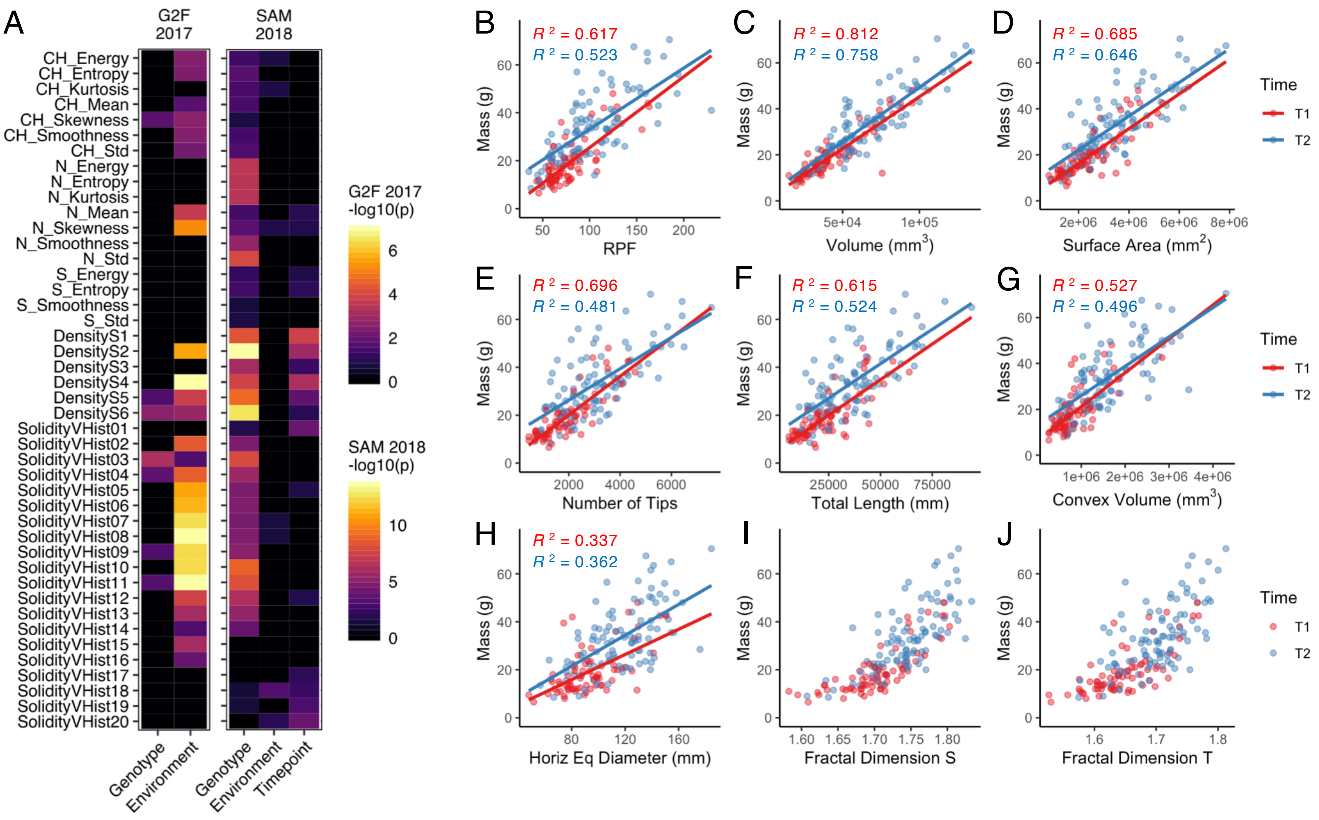

**Supplemental Figure 5:** Boxplots of values for all traits with significant differences (from ANOVA, adjusted p-values) between limited vs full irrigation in the G2F 2017 experiment.

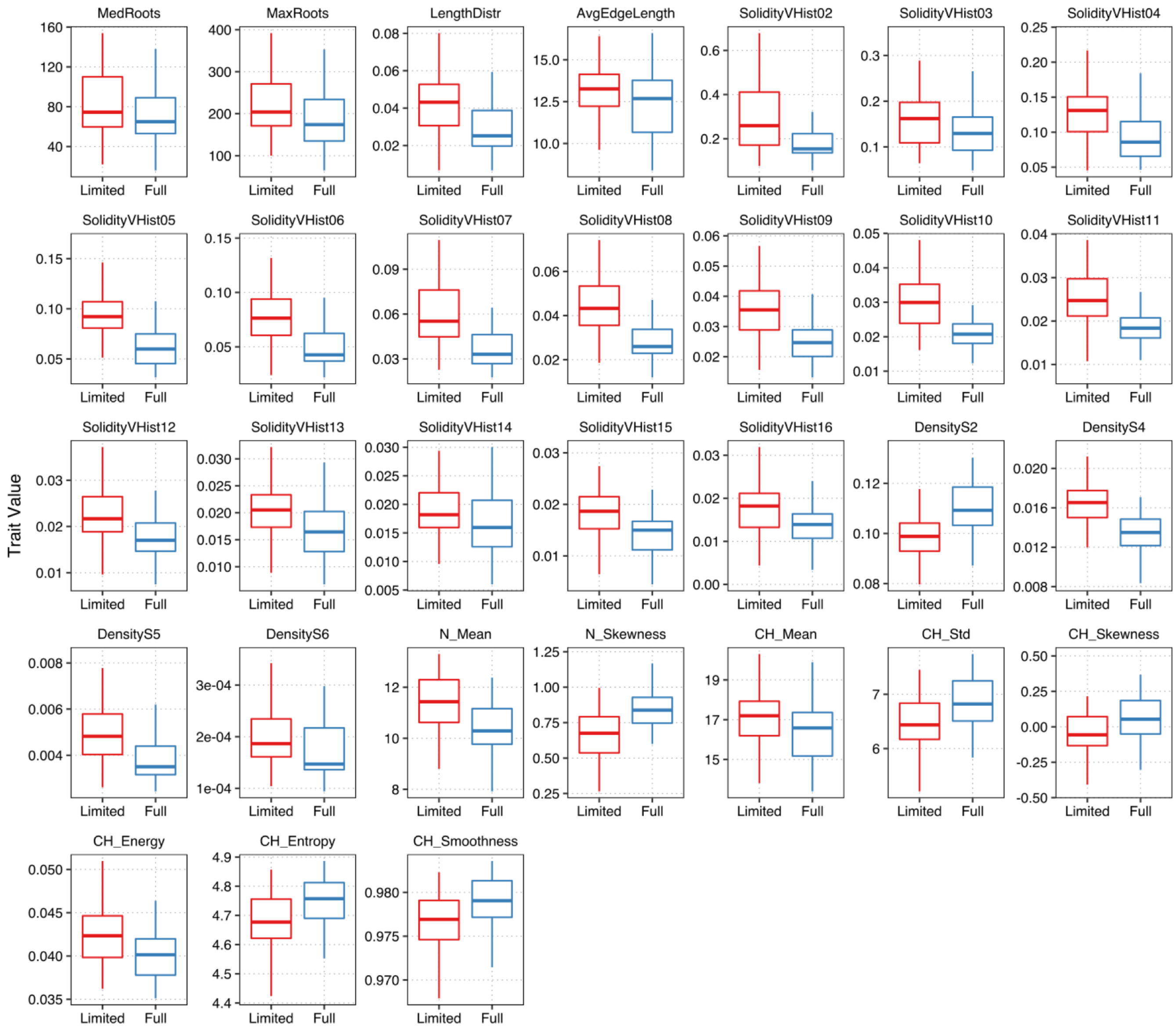

**Supplemental Figure 6:** Boxplots of values for all traits with significant differences (from ANOVA, adjusted p-values) between the 1st and 2nd time point or limited vs full irrigation in the SAM 2018 experiment.

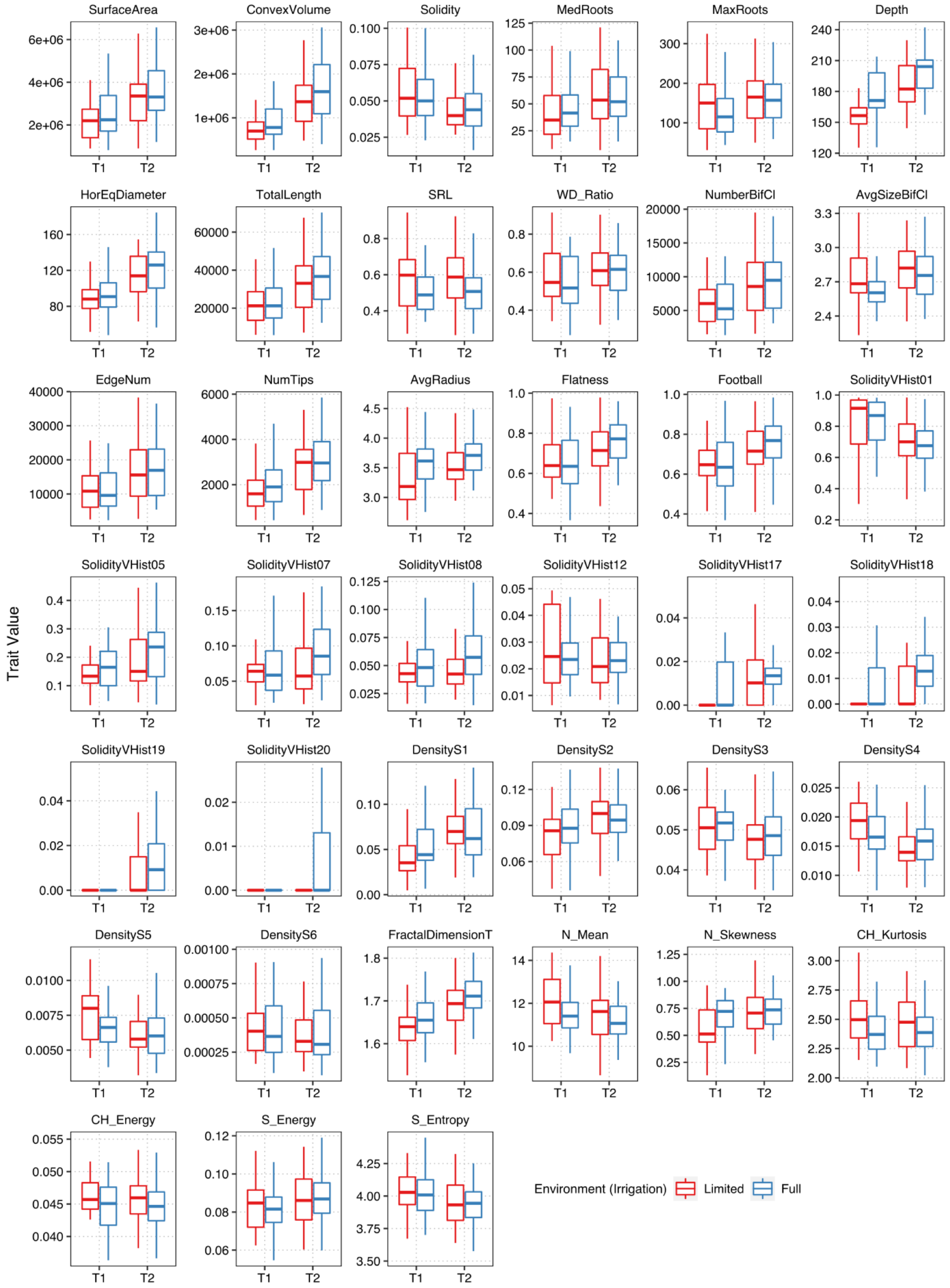

**Supplemental Figure 7:** Histogram of rank for every root trait across all PCA-LDA permutations (i.e., all possible combinations of 3 genotypes), in terms of importance for classification in the G2F 2017 data. Leftmost bin corresponds to 1st rank (better), while rightmost bin corresponds to 72nd rank (worse). Dashed line indicates rank expected by random chance if all traits were equal in importance.

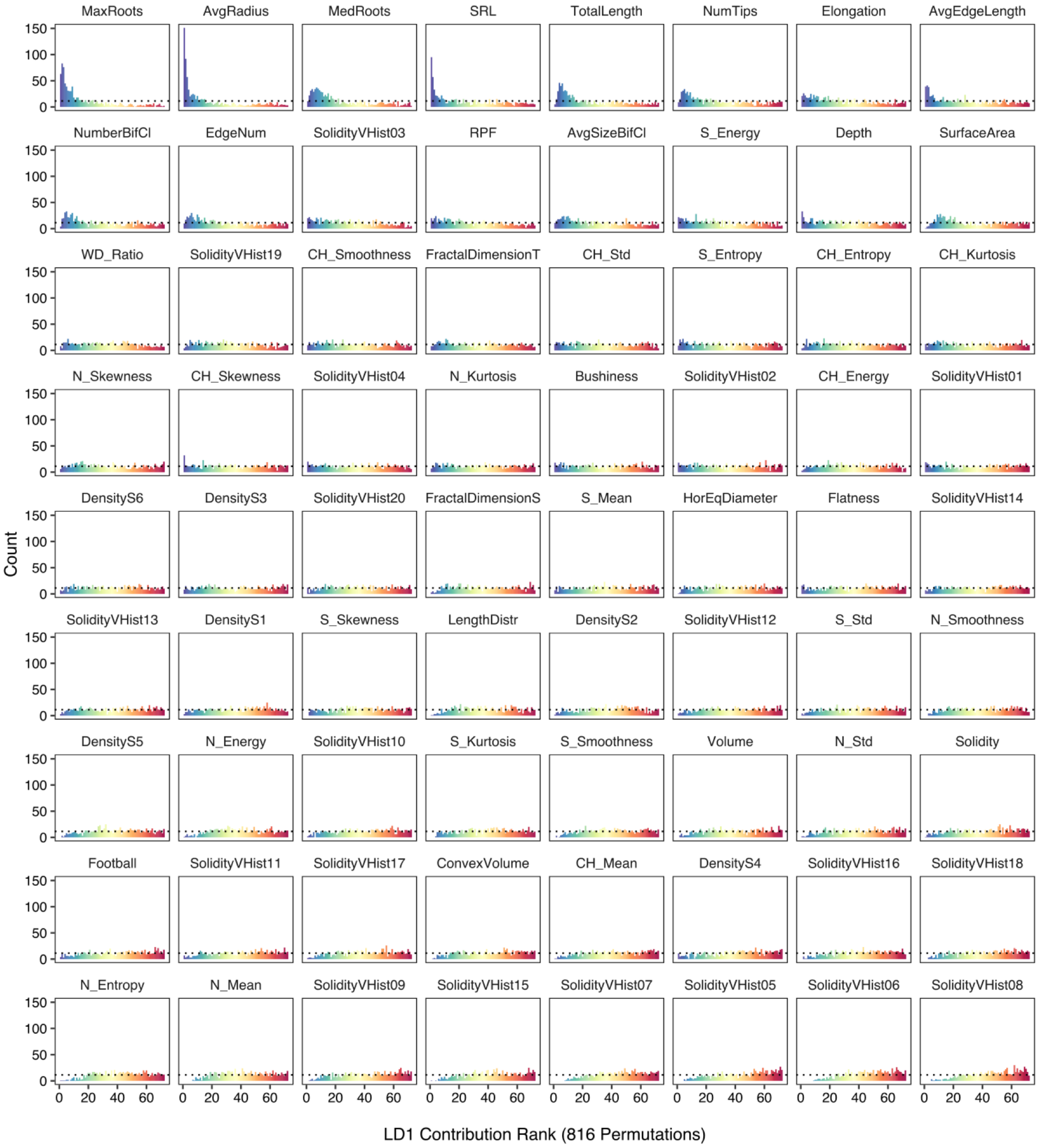

**Supplemental Figure 8:** Histogram of rank for every root trait across all PCA-LDA permutations (i.e., all possible combinations of 3 genotypes), in terms of importance for classification in the SAM 2018 data. Leftmost bin corresponds to 1st rank (better), while rightmost bin corresponds to 72nd rank (worse). Dashed line indicates rank expected by random chance if all traits were equal in importance.

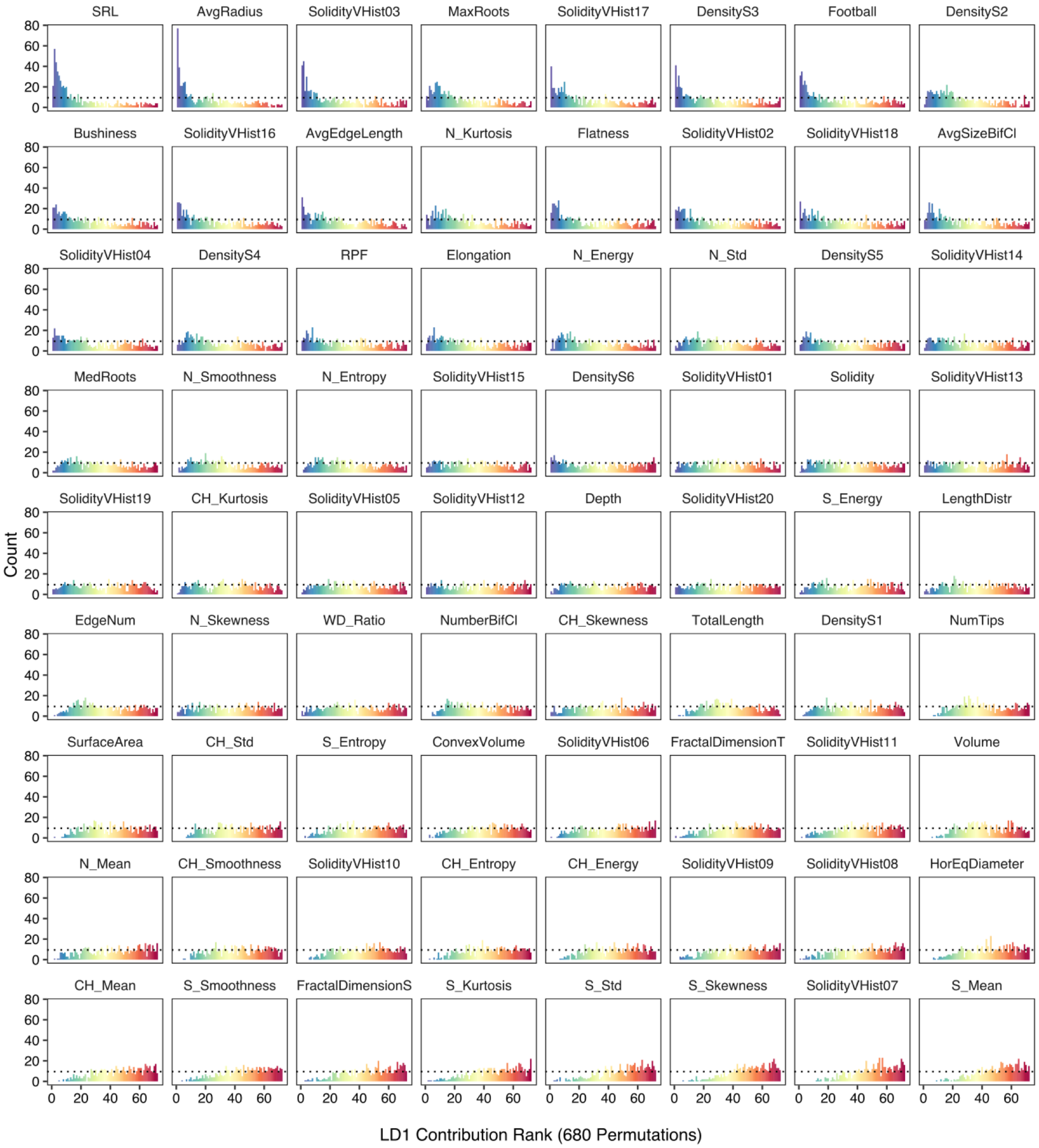

**Supplemental Figure 9:** (A) Visualization of PC1 and PC2 from principal component analysis of the G2M 2017 3D root data, colored by RPF value (note: the +/- direction of PC values is arbitrary). Linear regression between G2M 2017 PC2 values and RPF (B), and boxplot of between limited vs full irrigation environments for PC1 (C) and PC2 (D). (E) Visualization of PC1 and PC2 from principal component analysis of the SAM 2018 3D root data, colored by RPF value. Linear regression between SAM 2018 PC1 values and RPF (F), and boxplot of between limited vs full irrigation environments and time point 1 vs time point 2 for PC1 (C) and PC2 (D). 3D root trait loading estimates for PC1 and PC2 from 500 re-samplings (75% of dataset per re-sample) from G2M 2017 (I) and SAM 2018 (J) data.

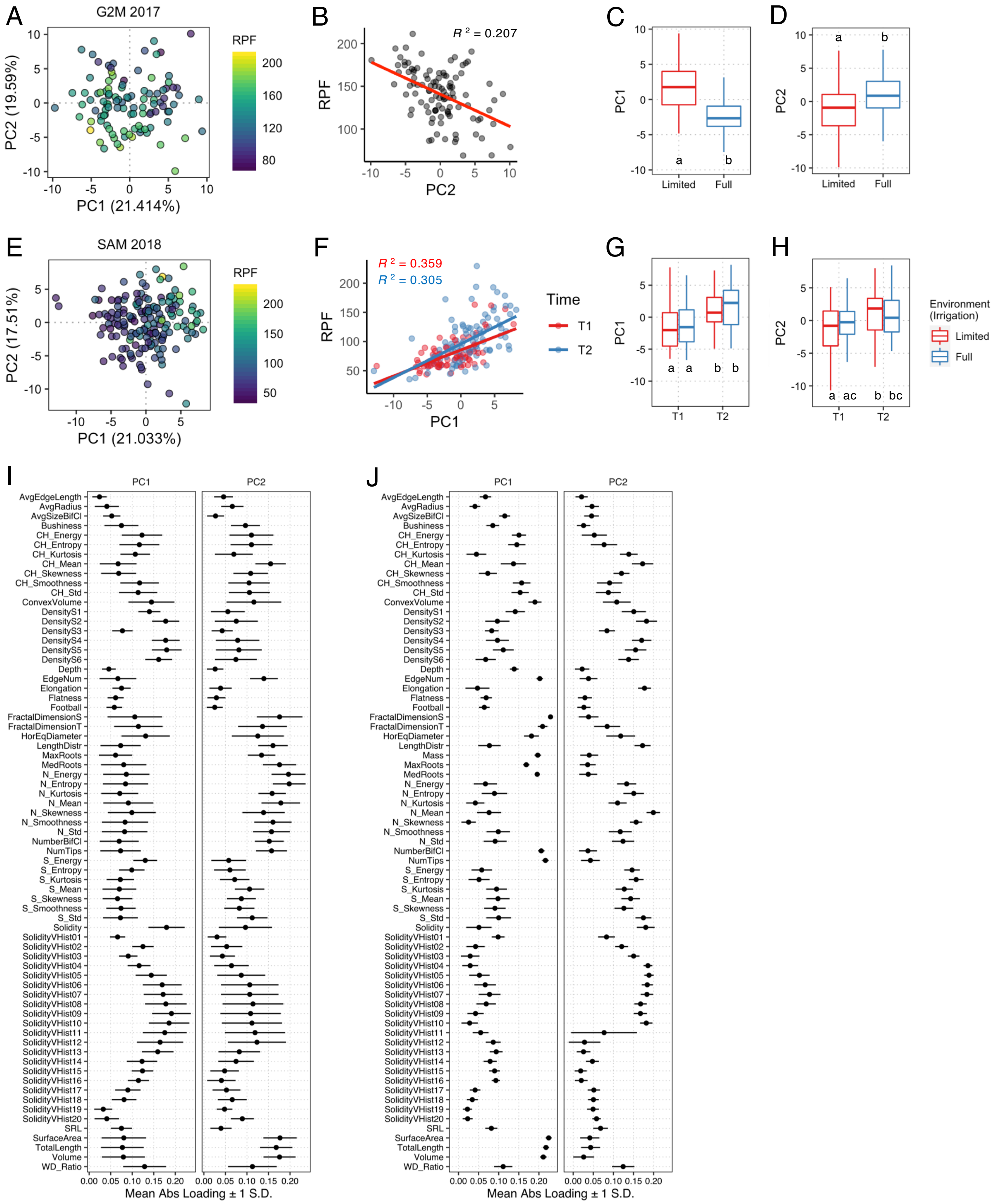
