## Supplemental Tables for "Complementary Phenotyping of Maize Root Architecture by Root Pulling Force and X-Ray Computed Tomography"

**Supplemental Table 1:** Genotypes included in 3D imaging from the G2M and SAM experiments.

| **G2M 2017** | **SAM 2018** |
| --- | --- |
| 15SFG:2148 | 778 |
| 15SJWE:G2F:04026/04020 | A441-5 |
| 15SJWE:G2F:05026/05020 | CH9 |
| 15SJWE:G2F:09041/09020 | CM7 |
| 15SJWE:G2F:12036/12020 | CR14 |
| 15SJWE:G2F:13041/13020 | H8431 |
| 15SJWE:G2F:14051/14020 | LH146Ht |
| 15SJWE:G2F:15046/15020 | M162W |
| 15SJWE:G2F:15051/15020 | Mt42 |
| 15SJWE:G2F:16031/16020 | N59B |
| 16SJWE:G2F:25251/25265 | NC236 |
| CS15-YCB-17-X-POL-J4 | Os420 |
| S15 IB-0364 | P39 |
| S15 IB-0384 | Pa875 |
| S16 CR-0369 | T8 |
| S16 CR-0438 | Tx601 |
| S16 CR-0462 | Va17 |
| S16 CR-0477 | Va99 |
| S16 CR-0485 | Z022E0104 |
| S16 CR-0511 | ZS01250 |
| S16 CR-0519 |  |
| S16 CR-0569 |  |
| S16 CR-0580 |  |
| S16 CR-0621 |  |
| S16 CR-0682 |  |
| S16 CR-0722 |  |
| S16 CR-0795 |  |
| S16 IA-0687 |  |
| WISN15/30649 |  |
| WISN15/30853 |  |

**Supplemental Table 2:** Traits descriptions; see Materials and Methods for citations to literature the traits are derived from, where applicable.

| **Trait** | **Description** |
| --- | --- |
| SurfaceArea | The sum of the local surface area at each voxel in the root network. |
| Volume | The sum of the local volume at each voxel in the root network; also referred to as “biomass.” |
| ConvexVolume | The volume of the convex hull that encompasses the root. |
| Solidity | The total network volume divided by the network convex volume. |
| MedRoots | Median number of roots among all horizontal slices. |
| MaxRoots | 84^th^ percentile value of the number of roots among all horizontal slices. |
| Depth | The number of voxels in the vertical direction of the root network. |
| HorEqDiameter | Maximum root network width. |
| TotalLength | Root length as approximated by the number of voxels in the network. |
| SRL | Specific root length; the total network length divided by the network volume. |
| LengthDistr | The ratio of root length in the upper 1⁄3 of the volume to the root length in the lower 2⁄3 of the volume. |
| WD_Ratio | Width-to-depth ratio; the maximum root network width divided by the network depth. |
| NumberBifCl | Number of bifurcation clusters. |
| AvgSizeBifCl | Average size of the bifurcation clusters. |
| EdgeNum | Number of skeleton edges. |
| AvgEdgeLength | Average length of skeleton clusters. |
| NumberTips | Number of root tips. |
| AvgRadius | The average radius of all voxels in the skeleton. |
| Elongation | PCA on 3D point cloud, taking the ratio between PC2 variance and PC1 variance; measures how elongated the root is. |
| Flatness | PCA on 3D point cloud, taking the ratio between PC3 variance and PC2 variance; measures how flat the root is. |
| Football | PCA on (x, y) of 3D point cloud, taking the ratio between PC2 variance and PC1 variance. |
| SolidityVHist 01-20 | The solidity at each slice is computed, then spline interpolated to the n^th^ cm (1-20) below the top. |
| DensityS 1-6 | The frequency of voxels with different 6 overlap ratios from side view. S6 represents the largest overlap ratio. Higher numbers in greater overlap ratio means a denser root. |
| FractalDimensionS | Fractal dimension is estimated from the projected side-view image using the box-counting method. It is a measure of how complicated a root shape is using self-similarity. |
| FractalDimensionT | Fractal dimension estimated from the projected top-view image using the box-counting method. It is a measure of how complicated a root shape is using self-similarity. |
| N/CH/S Mean | Mean estimated from the distribution of biomass/volume (N), convex hull (CH), or solidity (S) along the z-axis. |
| N/CH/S Std | Standard deviation estimated from the distribution of biomass/volume (N), convex hull (CH), or solidity (S) along the z-axis. |
| N/CH/S Skewness | Skewness, or inequality, estimated from the distribution of biomass/volume (N), convex hull (CH), or solidity (S) along the z-axis. Negative value indicates that a large number of the values are lower than the mean (left-tailed); positive value indicates that a larger number of the values are higher than the mean (right-tailed). |
| N/CH/S Kurtosis | Kurtosis, or peakiness, estimated from the distribution of biomass/volume (N), convex hull (CH), or solidity (S) along the z-axis. High value indicates that the peak of the distribution around the mean is sharp and long-tailed; low value indicates that the peak around the mean is round and short-tailed. |
| N/CH/S Energy | Energy, or uniformity, estimated from the distribution of biomass/volume (N), convex hull (CH), or solidity (S) along the z-axis. A high value indicates that the distribution has a small number of different levels. |
| N/CH/S Entropy | Entropy, the inverse of energy, estimated from the distribution of biomass/volume (N), convex hull (CH), or solidity (S) along the z-axis. A high value indicates that the distribution has a higher number of different levels. |
| N/CH/S Smoothness | Smoothness estimated from the distribution of biomass/volume (N), convex hull (CH), or solidity (S) along the z-axis. Defined as $1-\frac{1}{1+{(stddev)}^{2}}$ |

**Supplemental Table 3:** Classification accuracy summary from PCA-LDA (LOOCV) and random forest (10-fold CV). For PCA-LDA, sample size and number of groups (genotypes) per permutation is indicated. For random forest, number of groups (environments or time points) and sample size of the total model is indicated.

| **Data** | **Grouping** | **# Groups (Per Permutation)** | **Method** | **Sample Size  (Per Permutation)** | **Accuracy** |
| --- | --- | --- | --- | --- | --- |
| G2M 2017 | Genotype | 3 | PCA-LDA | 12 | 54.6% |
| G2M 2017 | Environment | 2 | Random Forest | 105 | 81.0% |
| SAM 2018 | Genotype | 3 | PCA-LDA | 27.75 (mean) | 67.2% |
| SAM 2018 | Environment | 2 | Random Forest | 171 | 66.2% |
| SAM 2018 | Time Point | 2 | Random Forest | 171 | 78.6% |

**Supplemental Table 4:** Spearman correlation between RPF and each 3D trait using every root sample as an observation. Adjusted p-values were calculated using the Benjamini-Hochberg method.

| **G2M 2017** | | | **SAM 2018** | | |
| --- | --- | --- | --- | --- | --- |
| **Trait** | **Correlation to RPF** | **Adj. p-value** | **Trait** | **Correlation to RPF** | **Adj. p-value** |
| FractalDimensionS | 0.64367077 | 9.25E-12 | Volume | 0.77528044 | 1.06E-33 |
| SurfaceArea | 0.61848397 | 7.31E-11 | FractalDimensionS | 0.749712 | 1.44E-30 |
| FractalDimensionT | 0.56966543 | 5.37E-09 | ConvexVolume | 0.74479439 | 3.95E-30 |
| Volume | 0.56082232 | 8.71E-09 | SurfaceArea | 0.70736033 | 5.26E-26 |
| TotalLength | 0.54212012 | 3.31E-08 | FractalDimensionT | 0.6883679 | 3.41E-24 |
| MedRoots | 0.53117402 | 6.58E-08 | HorEqDiameter | 0.66922067 | 1.72E-22 |
| ConvexVolume | 0.51922444 | 1.41E-07 | Depth | 0.64920791 | 7.84E-21 |
| HorEqDiameter | 0.50256974 | 4.16E-07 | TotalLength | 0.6260746 | 4.75E-19 |
| NumTips | 0.50041339 | 4.30E-07 | NumTips | 0.61795041 | 1.72E-18 |
| NumberBifCl | 0.43870857 | 2.03E-05 | NumberBifCl | 0.56105043 | 1.03E-14 |
| WD_Ratio | 0.41691289 | 5.73E-05 | SolidityVHist19 | 0.54799953 | 5.59E-14 |
| EdgeNum | 0.40390322 | 0.00010545 | EdgeNum | 0.54337232 | 9.49E-14 |
| MaxRoots | 0.38046053 | 0.00031647 | MedRoots | 0.46546205 | 7.71E-10 |
| N_Std | 0.36549113 | 0.00052685 | SolidityVHist18 | 0.45245647 | 2.64E-09 |
| N_Smoothness | 0.36549113 | 0.00052685 | DensityS1 | 0.44178551 | 6.91E-09 |
| CH_Entropy | 0.33243572 | 0.00209331 | SolidityVHist20 | 0.37134304 | 2.39E-06 |
| CH_Std | 0.31746048 | 0.00343406 | WD_Ratio | 0.33520294 | 2.77E-05 |
| CH_Smoothness | 0.31746048 | 0.00343406 | MaxRoots | 0.32494819 | 5.14E-05 |
| Elongation | 0.30162996 | 0.00597165 | SolidityVHist17 | 0.30719348 | 0.00014782 |
| N_Entropy | 0.29730689 | 0.00639042 | DensityS2 | 0.29980008 | 0.000219 |
| DensityS3 | 0.26218325 | 0.01748469 | CH_Std | 0.28322805 | 0.00049489 |
| S_Entropy | 0.25018855 | 0.02460862 | CH_Smoothness | 0.28322805 | 0.00049489 |
| SolidityVHist18 | 0.22670713 | 0.04446167 | SolidityVHist07 | 0.27461444 | 0.00075956 |
| AvgEdgeLength | 0.2223426 | 0.04868217 | CH_Entropy | 0.27089083 | 0.0008911 |
| SolidityVHist17 | 0.18297364 | 0.11532375 | SolidityVHist06 | 0.2490196 | 0.00250245 |
| SolidityVHist12 | 0.18170367 | 0.11575301 | SolidityVHist08 | 0.24575559 | 0.00282951 |
| SolidityVHist14 | 0.17664973 | 0.12680958 | Football | 0.24146799 | 0.00335381 |
| N_Mean | 0.15111563 | 0.19541674 | S_Energy | 0.23989478 | 0.00349689 |
| SolidityVHist16 | 0.14876748 | 0.20045463 | Flatness | 0.23852558 | 0.00355067 |
| LengthDistr | 0.14141796 | 0.22671434 | AvgRadius | 0.23362971 | 0.00426487 |
| DensityS2 | 0.14035461 | 0.22671434 | CH_Mean | 0.21543874 | 0.00893575 |
| SRL | 0.12237598 | 0.30954307 | AvgSizeBifCl | 0.18869802 | 0.02386921 |
| S_Std | 0.11599717 | 0.33223615 | N_Std | 0.17651548 | 0.03536019 |
| S_Smoothness | 0.11599717 | 0.33223615 | N_Smoothness | 0.17651548 | 0.03536019 |
| SolidityVHist15 | 0.11139937 | 0.3521347 | SolidityVHist05 | 0.17136987 | 0.04130921 |
| SolidityVHist20 | 0.10881279 | 0.35625924 | Elongation | 0.14747544 | 0.08372401 |
| SolidityVHist13 | 0.10841366 | 0.35625924 | SolidityVHist09 | 0.14662223 | 0.08409402 |
| S_Mean | 0.09929063 | 0.39313407 | N_Skewness | 0.13837582 | 0.09875453 |
| CH_Skewness | 0.08317502 | 0.48005671 | S_Kurtosis | 0.1221506 | 0.14539334 |
| AvgSizeBifCl | 0.06146119 | 0.61271982 | LengthDistr | 0.1069311 | 0.20416743 |
| SolidityVHist19 | 0.0464704 | 0.68948129 | N_Entropy | 0.09620175 | 0.24931259 |
| Solidity | 0.04604339 | 0.68948129 | S_Skewness | 0.09149535 | 0.27233255 |
| SolidityVHist11 | 0.01985294 | 0.85965405 | SolidityVHist04 | 0.08171773 | 0.32456168 |
| Depth | 0.0150373 | 0.87898932 | SolidityVHist10 | 0.07967893 | 0.33306625 |
| Football | -0.0189873 | 0.85965405 | N_Kurtosis | 0.03698406 | 0.66872693 |
| Flatness | -0.0305984 | 0.79004999 | SolidityVHist11 | 0.01752003 | 0.85626075 |
| SolidityVHist10 | -0.0363677 | 0.75518085 | SolidityVHist16 | 0.01251722 | 0.88598271 |
| CH_Mean | -0.0475486 | 0.68948129 | N_Mean | -0.0091104 | 0.90585945 |
| DensityS1 | -0.0551425 | 0.64955312 | SRL | -0.0122645 | 0.88598271 |
| S_Energy | -0.0612124 | 0.61271982 | AvgEdgeLength | -0.0563797 | 0.499047 |
| N_Skewness | -0.0775509 | 0.51081248 | N_Energy | -0.0583801 | 0.48954141 |
| DensityS4 | -0.092184 | 0.42800555 | SolidityVHist12 | -0.0824413 | 0.32456168 |
| SolidityVHist09 | -0.098876 | 0.39313407 | SolidityVHist03 | -0.101321 | 0.22538969 |
| SolidityVHist08 | -0.0989693 | 0.39313407 | SolidityVHist13 | -0.1058246 | 0.20607144 |
| S_Skewness | -0.1634628 | 0.15439143 | SolidityVHist02 | -0.1134866 | 0.17677503 |
| S_Kurtosis | -0.1704865 | 0.13550835 | SolidityVHist14 | -0.121763 | 0.14539334 |
| DensityS6 | -0.1737469 | 0.12898261 | S_Std | -0.1371182 | 0.09875453 |
| SolidityVHist07 | -0.1740994 | 0.12898261 | S_Smoothness | -0.1371182 | 0.09875453 |
| SolidityVHist05 | -0.1860423 | 0.11016656 | SolidityVHist15 | -0.1392362 | 0.09845089 |
| SolidityVHist01 | -0.1869235 | 0.11016656 | S_Mean | -0.1442558 | 0.08662018 |
| Bushiness | -0.2138888 | 0.05773223 | CH_Kurtosis | -0.1455866 | 0.08496011 |
| SolidityVHist04 | -0.2171124 | 0.05450343 | CH_Skewness | -0.151475 | 0.07567575 |
| SolidityVHist06 | -0.2285888 | 0.04352637 | DensityS3 | -0.1658295 | 0.04871094 |
| AvgRadius | -0.2401429 | 0.03220121 | DensityS6 | -0.1901895 | 0.0231517 |
| N_Kurtosis | -0.2671594 | 0.01543081 | Solidity | -0.1902681 | 0.0231517 |
| N_Energy | -0.274349 | 0.01262625 | S_Entropy | -0.2238772 | 0.00639915 |
| DensityS5 | -0.2765261 | 0.01220194 | SolidityVHist01 | -0.2382604 | 0.00355067 |
| SolidityVHist03 | -0.2864474 | 0.00902986 | Bushiness | -0.2658881 | 0.00111542 |
| SolidityVHist02 | -0.2994943 | 0.00616708 | CH_Energy | -0.2873189 | 0.00042849 |
| CH_Energy | -0.3765683 | 0.00035587 | DensityS4 | -0.358491 | 5.82E-06 |
| CH_Kurtosis | -0.4309385 | 2.87E-05 | DensityS5 | -0.4100719 | 1.13E-07 |

**Supplemental Table 5:** Spearman correlation between T1 and T2 for each trait using the average value for each genotype within time point as an observation.

| **Trait** | **T1-T2 Correlation** |
| --- | --- |
| DensityS5 | 0.894736842105263 |
| DensityS6 | 0.86015037593985 |
| Solidity | 0.86015037593985 |
| WD_Ratio | 0.846616541353383 |
| SRL | 0.837593984962406 |
| Bushiness | 0.793984962406015 |
| DensityS2 | 0.781954887218045 |
| Elongation | 0.768421052631579 |
| SolidityVHist03 | 0.748872180451128 |
| AvgSizeBifCl | 0.730827067669173 |
| HorEqDiameter | 0.726315789473684 |
| MedRoots | 0.724812030075188 |
| SolidityVHist05 | 0.715789473684211 |
| FractalDimensionT | 0.708270676691729 |
| DensityS1 | 0.705263157894737 |
| AvgEdgeLength | 0.685714285714286 |
| SolidityVHist04 | 0.682706766917293 |
| MaxRoots | 0.67218045112782 |
| DensityS4 | 0.669172932330827 |
| N_Std | 0.666165413533835 |
| SurfaceArea | 0.639097744360902 |
| N_Energy | 0.631578947368421 |
| FractalDimensionS | 0.631578947368421 |
| CH_Smoothness | 0.628571428571429 |
| N_Smoothness | 0.621052631578947 |
| N_Entropy | 0.621052631578947 |
| NumTips | 0.607518796992481 |
| AvgRadius | 0.593984962406015 |
| TotalLength | 0.590977443609023 |
| SolidityVHist06 | 0.584962406015038 |
| CH_Std | 0.580451127819549 |
| N_Kurtosis | 0.580451127819549 |
| EdgeNum | 0.56390977443609 |
| ConvexVolume | 0.556390977443609 |
| NumberBifCl | 0.538345864661654 |
| SolidityVHist09 | 0.521804511278195 |
| Flatness | 0.511278195488722 |
| SolidityVHist13 | 0.493233082706767 |
| CH_Entropy | 0.491729323308271 |
| RPF | 0.487719298245614 |
| Football | 0.463157894736842 |
| SolidityVHist10 | 0.44812030075188 |
| CH_Energy | 0.442105263157895 |
| CH_Mean | 0.427067669172932 |
| SolidityVHist12 | 0.42406015037594 |
| Volume | 0.416541353383459 |
| DensityS3 | 0.410526315789474 |
| CH_Skewness | 0.398496240601504 |
| SolidityVHist11 | 0.377443609022556 |
| Mass | 0.371929824561404 |
| N_Mean | 0.369924812030075 |
| SolidityVHist07 | 0.356390977443609 |
| S_Energy | 0.353383458646617 |
| S_Entropy | 0.33984962406015 |
| SolidityVHist08 | 0.332330827067669 |
| LengthDistr | 0.320300751879699 |
| SolidityVHist02 | 0.308270676691729 |
| S_Smoothness | 0.303759398496241 |
| SolidityVHist14 | 0.270676691729323 |
| N_Skewness | 0.266165413533835 |
| CH_Kurtosis | 0.239097744360902 |
| S_Mean | 0.237593984962406 |
| Depth | 0.227067669172932 |
| S_Skewness | 0.198496240601504 |
| S_Kurtosis | 0.172932330827068 |
| S_Std | 0.16390977443609 |
| SolidityVHist01 | 0.103759398496241 |
| SolidityVHist15 | -0.00902255639097744 |
| SolidityVHist18 | -0.0360902255639098 |
| SolidityVHist16 | -0.159398496240601 |
| SolidityVHist17 | -0.160902255639098 |
| SolidityVHist19 | -0.279699248120301 |
| SolidityVHist20 | -0.539157237939949 |
